# The SARS-CoV-2 reproduction number *R_0_* in cats

**DOI:** 10.1101/2021.07.20.453027

**Authors:** Jose L. Gonzales, Mart C.M. de Jong, Nora M. Gerhards, Wim H. M. Van der Poel

## Abstract

Domestic cats are susceptible to SARS-CoV-2 virus infection and given that they are in close contact with people, assessing the potential risk cats represent for the transmission and maintenance of SARS-CoV-2 is important. Assessing this risk implies quantifying transmission from humans-to-cats, from cats-to-cats and from cats-to-humans. Here we quantified the risk of cat-to-cat transmission by reviewing published literature describing transmission either experimentally or under natural conditions in infected households. Data from these studies were collated to quantify the SARS-CoV-2 reproduction number *R_0_* among cats. The estimated *R_0_* was significantly higher than 1, hence cats could play a role in the transmission and maintenance of SARS-CoV-2. Questions that remain to be addressed are the risk of transmission from humans-to-cats and cats-to-humans. Further data on household transmission and data on virus levels in both the environment around infected cats and their exhaled air could be a step towards assessing these risks.

---

A relevant concern in the control of the ongoing Covid-19 pandemic is the risk domestic animals could play in the maintenance and transmission of SARS-CoV-2. Assessing this risk implies quantifying transmission from humans-to-animals, from animals-to-animals and from animals-to-humans. Large epidemics in farmed minks have confirmed this risk for that specific species (1). The role of cats is of particular interest, because they are in close contact with humans and frequently in contact with other cats. Available field (2–9) and experimental data (10–14) indicate that cats are susceptible to infection, occasionally show mild clinical signs and may be able to transmit the infection between cats. Indeed, transmission experiments confirmed this possibility (10–14), however, the lack of a proper statistical assessment of transmission in the reported experiments limits confident extrapolation of the results from the experiment to the population. An important question when assessing the risk of transmission is whether cat-to-cat transmission can be sustained. A key measure to answer this question is the basic reproduction number *R_0_*, which is the average number of individuals to whom a typical infectious individual will transmit the infection to in a naive population. *R_0_* is a key parameter in infectious disease epidemiology, it provides an indication of the transmissibility of a pathogen and the risk of epidemic transmission. When *R_0_* > 1, one can expect sustained transmission with high risk of a major outbreak and endemicity to occur, whereas when *R_0_* < 1 the infection is likely to peter out. Other parameters which contribute to quantitatively describe transmission are: 1) the latent period *L,* which is the time from becoming infected to becoming contagious, 2) the infectious period *T,* which is the average period of time an individual is contagious and 3) the transmission rate parameter *β* which is the number of contact infections caused by one typical infectious individual per unit of time. Here, published experiments and observational studies describing infection and transmission of SARS-CoV-2 between cats were reviewed. Data from these studies were collated and analysed to statistically confirm whether cat-to-cat transmission can be sustained and to provide estimates of relevant transmission parameters.

A systematic literature search was conducted which identified 115 publications. Upon screening and selection of relevant studies for data collection and analysis, five experimental studies and 8 observational studies were included for analysis. A detailed description of the systematic review process is provided as supplemental material (Text S1).

In Tables 1 and 2 the experimental and household studies included for analyses are summarised. Of the experimental studies, four (10–12) assessed direct-contact transmission and one (13) indirect (droplet) transmission. These studies used different study designs with respect to age and the number of inoculated (donor) and contact cats included within an experimental group. All experiments used inoculation doses ≥ 10^5^ PFU (Gaudreault et al (12) used 10^6^ TCID_50_) and the predominant inoculation route was intra-nasal inoculation. Following inoculation, infection and transmission were monitored by longitudinally detecting and measuring virus shedding in nasal, faecal or oropharyngeal samples collected from inoculated and contact- or droplet-infected cats. The laboratory methods used to monitor infection were either virus isolation (VI) (10, 11) or RT-PCR (12, 13). From the observational studies, data from 12 households housing infected people and at least one infected cat were included for analysis. Eight of these households (4–9) had either two or three cats and four households (15, 16) had only one cat (Tables 2, S3, S4). The infection process of owners and cats was longitudinally followed in most of these households.

**Table 1.** Summary of the experimental procedures showing the study design, the age of the cats, the inoculation route and dose, the type of samples taken and the diagnostic method used to quantify virus levels in time.

| *Study* | Type of transmission | Design I x S<sup>a</sup> | Cat's age (months) | Inoculation Route | Dose (log<sub>10</sub>) | Units<sup>b</sup> | Sample (route)<sup>c</sup> | Diagnostic test |
| --- | --- | --- | --- | --- | --- | --- | --- | --- |
| Halfmann et al.(10) | Direct contact | 1 x 1 | 3.5 to 4.2 | Nasal,Tracheal, Oral,Ocular | 5.7 | PFU | Respiratory | VI<sup>d</sup> |
| Bosco-Lauth et al.(11) | Direct contact | 2 x 2 | 60 - 96 | Nasal | 5.4 | PFU | Respiratory/rectal | VI |
| Gaudreault et al. (12) | Direct contact | 3 x 1 | 4.5 - 5 | Nasal, Oral | 6 | TCID<sub>50</sub> | Respiratory | RT-PCR |
| Bao et al.(14) | Direct contact | 1 x 1 | 8 - 18 | Nasal | 6 | TCID<sub>50</sub> | Respiratory/rectal | RT-PCR |
| Shi et al. juveniles.(13) | Indirect-droplet | 1 x 1 | 2.3 to 3.3 | Nasal | 5 | PFU | Respiratory | RT-PCR |
| Shi et al. subadults.(13) | Indirect-droplet | 1 x 1 | 6 to 9 | Nasal | 5 | PFU | Rectal | RT-PCR |

<sup>a</sup> I = number of inoculated cats and S = number of susceptible contacts per group at the start of the experiment.
<sup>b</sup> PFU = Plaque-forming units, TCID<sub>50</sub> = Fifty-percent tissue culture infective dose
<sup>c</sup> Type of samples considered as respiratory were: nasal swabs, oropharyngeal swabs, nasal washes. Rectal samples were: rectal swabs or faeces.
<sup>d</sup> VI = Virus Isolation

**Table 2.** Summary description of the households studies included for estimation of the shedding (infectious) period and the Reproductive Number *R_0_.*

| Studies | No. of households | Total No. of cats per household | Number of households with > 1 cat infected<sup>a</sup> | Sample (Route)<sup>b</sup> | Diagnostic test | Data used for estimation of |
| --- | --- | --- | --- | --- | --- | --- |
| Chaintoutis et al. (4), Hamer et al.(7), Neira et al.(6) | 3 | 3 | 1 | Respiratory/rectal | PCR, Serology | $R_0$, Shedding |
| Hamer et al.(7), Klaus et al.(5), Segales et al.(9), Neira et al.(6) Goryoka et al.(8) | 5 | 2 | 3 | Respiratory/rectal | PCR, Serology | $R_0$, Shedding |
| Barrs et al.(15), Bessiere et al.(16) | 4 | 1 |  | Respiratory/rectal | PCR, Serology | Shedding |

<sup>a</sup> For a cat to be considered infected it had to be seropositive the last time the cats in the household were sampled.
<sup>b</sup> Type of samples considered as respiratory were: nasal swabs, oropharyngeal swabs or oral swabs. Rectal samples were: rectal swabs or faeces.

For the statistical analysis of the transmission experiments, temporal data on infection of inoculated and contact-or droplet-infected cats was collected. Within each experimental group, an inoculated cat was classed as infectious when it was reported as shedding virus, regardless of the viral load and of the detection method (virus isolation or RT-PCR). Contact cats were considered susceptible for the period of days before the first day they were shown to shed virus (one day latent period (Table 3)). The prepared datasets (Tables S1, S2) were used to estimate *L (days)*, *T (days)*, **β* (day^−1^)* and *R_0_*. The first two parameters were estimated using parametric survival regression models, *β* was estimated by using a SEIR model fitted by using a generalised linear regression model and *R_0_* was estimated either as the product of *T* * *β* or by using the final size method (FSM). The latter only requires information of the total number of infections in a group/household at the end of the infection process, when there is either no more infectious or no more susceptible hosts present (17, 18). For analysis of the household data (Table 2), transmission was analysed using the FSM, and the length of shedding was estimated using parametric survival models. To simplify the analysis of transmission, it was assumed that the source of infection of secondary infected cats was the first infected cat (infected by the owner) in the household and the contribution of infected owners to the infection of secondary infected cats was not included in the analysis. A detailed explanation of the statistical analysis is provided as Supplemental material (Text S2).

**Table 3.**
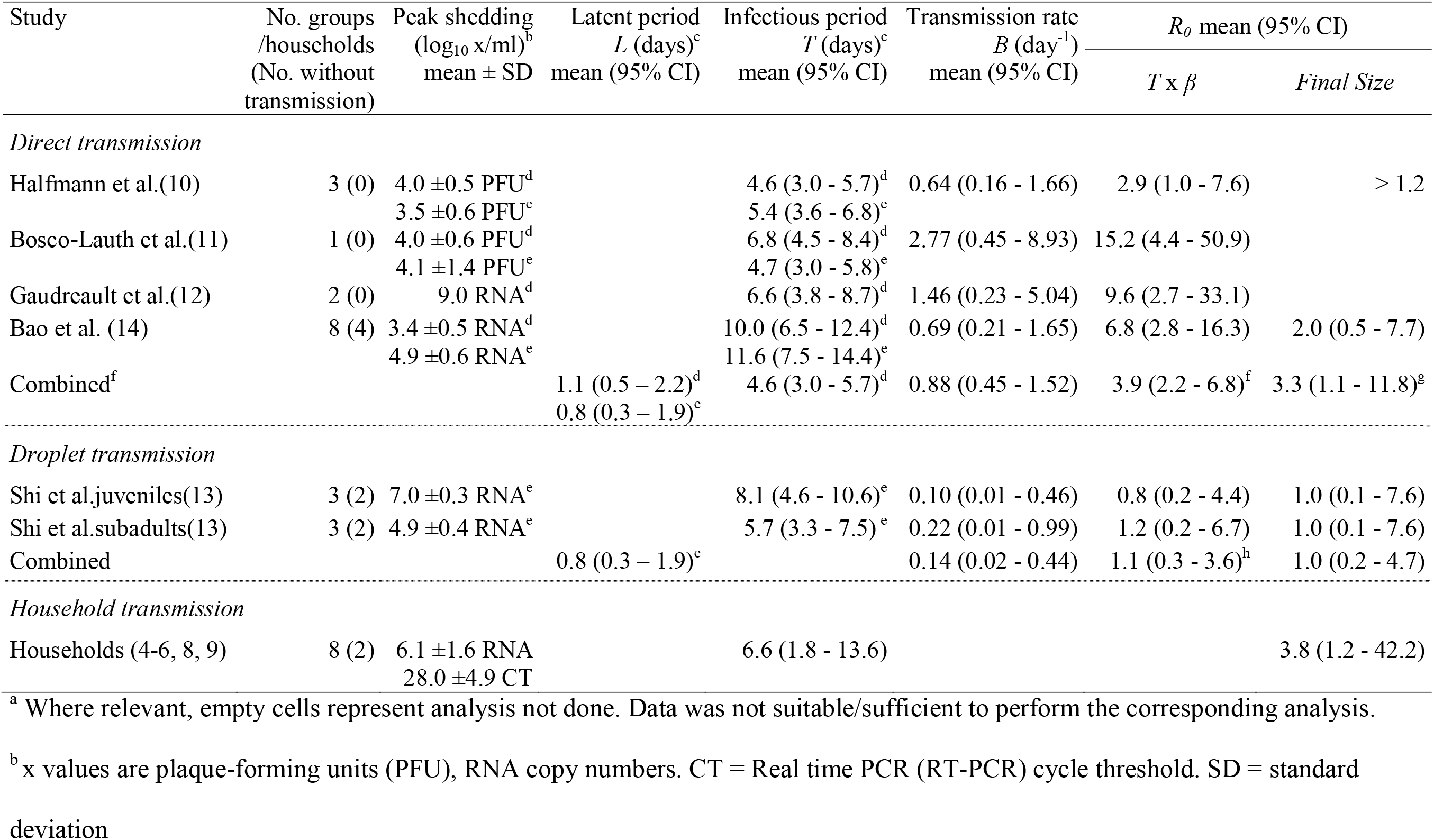
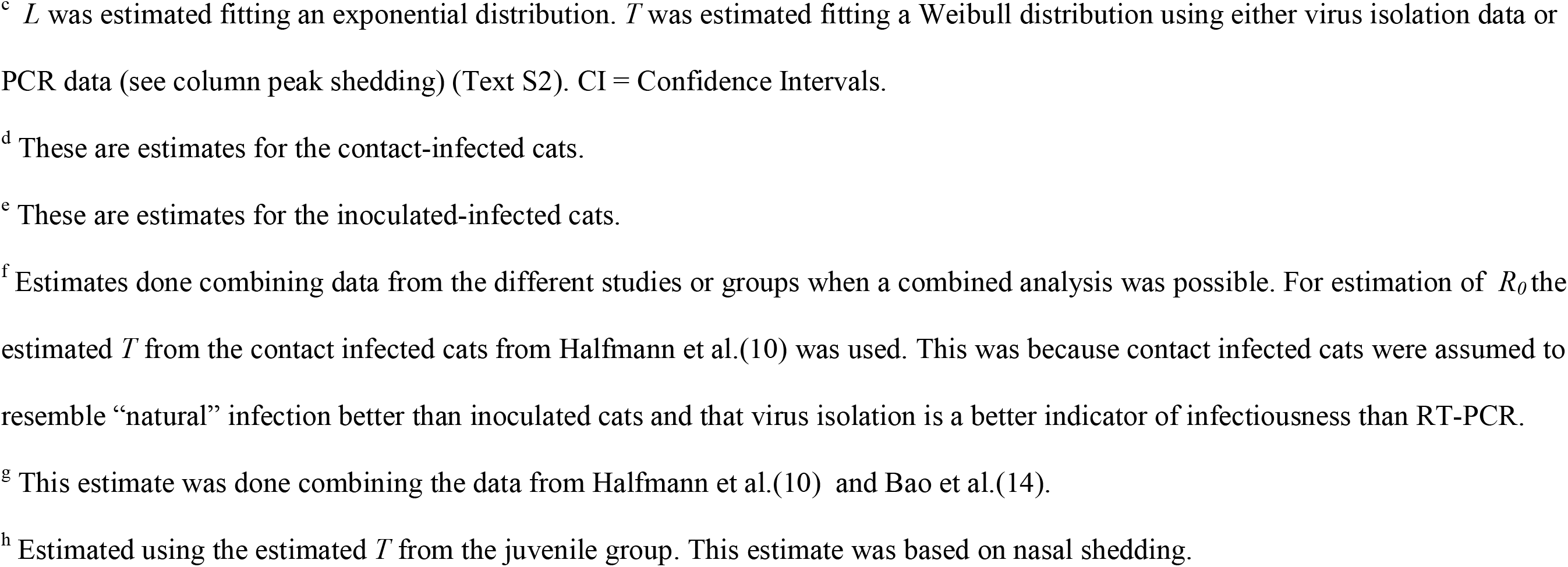
Quantified parameters for direct contact and droplet transmission of SARS-CoV-2 between cats using data from transmission experiments or observational studies describing infection and transmission at household level.^a^

| Study | No. groups<br>/households<br>(No. without<br>transmission) | Peak shedding<br>(log<sub>10</sub> x/ml)<sup>b</sup><br>mean ± SD | Latent period<br>*L* (days)<sup>c</sup><br>mean (95% CI) | Infectious period<br>*T* (days)<sup>c</sup><br>mean (95% CI) | Transmission rate<br>*B* (day<sup>-1</sup>)<br>mean (95% CI) | *R*<sub>0</sub> mean (95% CI) |  |
| --- | --- | --- | --- | --- | --- | --- | --- |
|  |  |  |  |  |  | *T* x *β* | *Final Size* |
| *Direct transmission* |  |  |  |  |  |  |  |
| Halfmann et al.(10) | 3 (0) | 4.0 ±0.5 PFU<sup>d</sup><br>3.5 ±0.6 PFU<sup>e</sup> |  | 4.6 (3.0 - 5.7)<sup>d</sup><br>5.4 (3.6 - 6.8)<sup>e</sup> | 0.64 (0.16 - 1.66) | 2.9 (1.0 - 7.6) | > 1.2 |
| Bosco-Lauth et al.(11) | 1 (0) | 4.0 ±0.6 PFU<sup>d</sup><br>4.1 ±1.4 PFU<sup>e</sup> |  | 6.8 (4.5 - 8.4)<sup>d</sup><br>4.7 (3.0 - 5.8)<sup>e</sup> | 2.77 (0.45 - 8.93) | 15.2 (4.4 - 50.9) |  |
| Gaudreault et al.(12) | 2 (0) | 9.0 RNA<sup>d</sup> |  | 6.6 (3.8 - 8.7)<sup>d</sup> | 1.46 (0.23 - 5.04) | 9.6 (2.7 - 33.1) |  |
| Bao et al. (14) | 8 (4) | 3.4 ±0.5 RNA<sup>d</sup><br>4.9 ±0.6 RNA<sup>e</sup> |  | 10.0 (6.5 - 12.4)<sup>d</sup><br>11.6 (7.5 - 14.4)<sup>e</sup> | 0.69 (0.21 - 1.65) | 6.8 (2.8 - 16.3) | 2.0 (0.5 - 7.7) |
| Combined<sup>f</sup> |  |  | 1.1 (0.5 – 2.2)<sup>d</sup><br>0.8 (0.3 – 1.9)<sup>e</sup> | 4.6 (3.0 - 5.7)<sup>d</sup> | 0.88 (0.45 - 1.52) | 3.9 (2.2 - 6.8)<sup>f</sup> | 3.3 (1.1 - 11.8)<sup>g</sup> |
| *Droplet transmission* |  |  |  |  |  |  |  |
| Shi et al.juveniles(13) | 3 (2) | 7.0 ±0.3 RNA<sup>e</sup> |  | 8.1 (4.6 - 10.6)<sup>e</sup> | 0.10 (0.01 - 0.46) | 0.8 (0.2 - 4.4) | 1.0 (0.1 - 7.6) |
| Shi et al.subadults(13) | 3 (2) | 4.9 ±0.4 RNA<sup>e</sup> |  | 5.7 (3.3 - 7.5)<sup>e</sup> | 0.22 (0.01 - 0.99) | 1.2 (0.2 - 6.7) | 1.0 (0.1 - 7.6) |
| Combined |  |  | 0.8 (0.3 – 1.9)<sup>e</sup> |  | 0.14 (0.02 - 0.44) | 1.1 (0.3 - 3.6)<sup>h</sup> | 1.0 (0.2 - 4.7) |
| *Household transmission* |  |  |  |  |  |  |  |
| Households (4-6, 8, 9) | 8 (2) | 6.1 ±1.6 RNA<br>28.0 ±4.9 CT |  | 6.6 (1.8 - 13.6) |  |  | 3.8 (1.2 - 42.2) |

<sup>a</sup> Where relevant, empty cells represent analysis not done. Data was not suitable/sufficient to perform the corresponding analysis.
<sup>b</sup> x values are plaque-forming units (PFU), RNA copy numbers. CT = Real time PCR (RT-PCR) cycle threshold. SD = standard deviation
<sup>c</sup> $L$ was estimated fitting an exponential distribution. $T$ was estimated fitting a Weibull distribution using either virus isolation data or PCR data (see column peak shedding) (Text S2). CI = Confidence Intervals.
<sup>d</sup> These are estimates for the contact-infected cats.
<sup>e</sup> These are estimates for the inoculated-infected cats.
<sup>f</sup> Estimates done combining data from the different studies or groups when a combined analysis was possible. For estimation of $R_0$ the estimated $T$ from the contact infected cats from Halfmann et al.(10) was used. This was because contact infected cats were assumed to resemble “natural” infection better than inoculated cats and that virus isolation is a better indicator of infectiousness than RT-PCR.
<sup>g</sup> This estimate was done combining the data from Halfmann et al.(10) and Bao et al.(14).
<sup>h</sup> Estimated using the estimated $T$ from the juvenile group. This estimate was based on nasal shedding.

For all experiments, *L* was estimated to be about one day, with no significant differences observed between inoculated and contact infected cats (Table 3). The type of test has a clear influence in the estimation of *T*, with estimates done using RT-PCR data leading to an overestimation of *T* and consequently *R_0_* when compared with the FSM estimates. Using VI data from contact-infected cats (assumed to closely reflect a “natural” infection) to estimate *T* and the corresponding *R_0_* led to similar estimates to those done using the FSM (Table 2). The experimental design had a large influence in the estimation of *β*; with the design used in two of the studies (11, 12) leading to an overestimation of this parameter and large standard errors. Although a small sample size was used, the pair-transmission design used by Shi et al (13), Halfmann et al (10) and Bao et al. (14) allowed the estimation of *β* and *R_0_* with good certainty. The former experiment assessed droplet-transmission whilst the latter two experiments assessed direct transmission and allowed confirmation that *R_0_* is significantly higher than 1 (p < 0.05). When combining these two experiments, the estimated *R_0_* (*T* * *β*) for cats was 3.9 (95% confidence intervals: 2.2 – 6.8) or 3.3 (FSM) (1.1 – 11.8). These estimates were similar to the estimates done at household level, with the estimated *R_0_* (FSM) being 3.8 (1.2 – 42.2) (Table 3). Similarly, the estimates of *T* and virus shedding levels from household data were similar to those estimates from the experiments (Table 3). Noting the assumptions made for the analysis of household data, the results indicate that pair-transmission experiments appear to provide a reliable approximation of the expected transmission dynamics of SARS-CoV-2 between cats at household level. Compared to direct transmission, droplet transmission was slower *β =* 0.14 (0.02 – 0.44) *day ^−1^* and may happen to a lower extend *R_0_* = 1.0 (0.2 – 4.7) than direct transmission (Table 3).

This study shows the importance of quantitatively assessing transmission when performing transmission experiments and the relevance of a proper experimental design to obtain reliable estimates of different parameters that describe the transmission process. Pair-transmission experiments are a suitable design to assess transmission. By using both data from the studies that used this type of experimental design (10, 14) and data from studies which followed infected households, we statistically confirmed that sustained transmission of SARS-CoV-2 among cats can be expected (*R_0_* > 1). To put this into perspective, scenarios in which contacts between stray and household cats take place (3) could lead to persistence of the virus in the cat population.

By combining field and experimental observations we could partly validate the suitability of pair-transmission experiments to study transmission and the validity of the estimated parameters. Whilst field observations would be ideal, it is practically impossible to obtain detailed temporal data to have a thorough understanding of the transmission dynamics. Given this limitation, in order to analyse the household data we had to make assumptions which influence our estimates. The main assumption being that secondary infected cats were infected by the first infected cat in the household, ignoring the possibility of these cats becoming infected by contact with the infected owner. As a result the *R_0_* estimates could be overestimated. As for *T* and shedding levels, observations were left censored, since first diagnosis of the cats was around five to seven days after clinical onset of the infected owner (Table S4) and not all cats were followed daily, which may affect the accuracy of these estimates. Nevertheless, they were similar to the experimental estimates. The combination of experimental and field data in this study improved the characterization of transmission between cats and increased the certainty in the estimated parameters.

Interestingly, levels of virus shedding in household infected cats, were as high as those observed experimentally (Table 3), with reported shedding levels as high as 10^8.5^ RNA copies/swab sample or RT-PCR CT values as low as 21 (Table S4). Considering both that infected cats shed high levels of virus, and that droplet transmission is possible, the risk for cat-to-human transmission of SARS-CoV-2 may not be low. There is a need to further investigate this risk. Experimental assessment of, for example, the probability of transmission via a contaminated environment around an infected cat and measurements of virus concentrations in infected cats’ exhaled air would provide further information to quantify the risk for cat-to-human transmission. This data combined with more detailed transmission and environmental contamination data (5, 16) from infected household cats could aid to further quantify the combined risks of human-to-cat and cat-to-human transmission. Thorough understanding of transmission of SARS-CoV-2 at the human-animal interplay is important to obtain a better insight into the population dynamics of this virus.

## Acknowledgments

This work was supported by funding from the European Union’s Horizon 2020 Research and Innovation programme under grant agreement No 773830: One Health European Joint Programme, projects MATRIX and COVRIN. We thank Michel Counotte for his help with the literature search.

No potential conflict of interest was reported by the author(s)

## Supplemental material

**Text S1.** Literature search and selection of manuscripts

**Text S2.** Data analysis methods for the estimation of transmission parameters

**Table S1.** Collated data for the quantification of the transmission rate *β* (day^−1^). Data for each pair of cats (inoculated + contact) was collated daily from day one post inoculation to the day the contact cat was assumed infected (one day before shedding virus).

**Table S2.** Collated data for the estimation of the infectious and latent periods.

**Table S3.** Collated data from infected households with more than one cat. These data were used for the estimation of the reproductive number R_0_ using the final size method.

**Table S4.** Collated data from observational studies describing the longitudinal follow up of infection in infected cats from infected households. These data were used to estimate the duration of observed shedding in naturally infected cats.

**Table S5.** Estimated Weibull parameters (Shape and Scale) describing the length of the infectious period *T.*

