## Supplemental Text S1 for "The SARS-CoV-2 reproduction number *R_0_* in cats"

### Text S1. Literature search and selection of manuscripts

*Search question*

For our research question we defined the study population as naive domestic cats exposed, experimentally or naturally within a household, to SARS-CoV-2. The intervention of interest was infection via experimental challenge or household infection and the outcomes of interest were cat-to-cat transmission and longitudinal monitoring of shedding in infected cats.

*Search strategy*

We searched the electronic databases PubMed, EMBASE, BioRxiv, and MedRxiv as indexed by the 'COVID Open Access Project'(1). We filtered the dataset of COVID-19 related research based on keywords in the title and or abstract: "((cats) OR (feline)) AND ((transmission) OR (transmissibility) OR (passage) OR (susceptibility))". The search was performed on 10 May 2021. Additionally, we checked the reference lists of the included studies for relevant publications.

*Study selection and data extraction*

First screening and selection was made based on titles and abstracts. Only experimental studies and observational studies in line with the research question were included. The second step consisted of full text screening of the selected studies. For experimental studies, studies which included naive sentinels exposed directly or indirectly to infected (inoculated) cats to assess transmission were selected for data extraction. For observational studies, selected studies for data extraction were reports in which detailed information of infection of household cats were provided at household level. Studies with one cat or multiple cats per household were selected for data extraction.

Data was extracted either from the text, figures or the supplementary data files. For studies that did not provide the full data, we used WebPlotDigitizer (2) to extract the data from the figures. Data on the first and last time of positive RNA or virus detection during the infection process, peak virus concentration (units depended on the laboratory method used), the number of cats per experimental group or household and the number infected cats at the end of the study were extracted for analysis. Detailed description of data extraction and preparation is provided in Text S2.

*Literature search and study selection results*

The literature search provided 115 studies (111 identified for the searched databases and four by checking reference lists). Twenty nine (3-31) studies were then selected for full text screening and are listed below. Finally 13 studies, five experimental studies (4, 8, 14-16) and eight observational studies (5, 6, 10, 17-20, 31) were selected for data extraction and analysis.
