## Supplemental Text S2 and Tables for "The SARS-CoV-2 reproduction number *R_0_* in cats"

### Text S2. Data analysis methods for the estimation of transmission parameters

#### *Data preparation*

*Transmission experiments*

Daily presence of infection in individual cats as measured by virus shedding in nasal (swabs or washes) (1-5) or faecal samples (4) from inoculated and contact infected cats was extracted from the published articles. Within each experimental group (inoculated + contact) a cat was classed as infectious when it was shown as shedding virus, regardless of the viral load. Contact cats were considered susceptible for the period of days before the first day (after estimation of the latent period) they were shown to shed virus. For data collation a latent period (time from infection to becoming infectious) of one day was used. One day was used because the latent period was estimated to be 1.1 (95% Confidence intervals: 0.5 – 2.2) days for contact=infected cats and 0.84 (0.5 – 1.4) days for inoculated cats, with no significant difference observed in the latent period between contact-infected and inoculated cats (Table 3, main text). Table S1 shows how this data were prepared.

For estimation of the mean peak of shedding, the highest virus load recorded for each animal during the experiment period was extracted for analysis. Data preparation for the estimation of the latent and infectious period is explained below and shown in Table S2.

*Household observations (Observational studies)*

For the assessment of transmission, data from households housing more than one cat were selected. From the selected households, the following data were extracted: number of people living in the household (at least one person infected), the total number of cats in the household, the number of cats diagnosed as positive by RT-PCR and serology as well as the last day that the cat was tested for serology (Table S3).

For the estimation of the peak and length of shedding (Infectious period), data from households housing one or more infected cats which were longitudinally followed after first diagnosis were extracted for analysis. Table S4 shows how these data were extracted and prepared for analysis.

#### *Estimation of the transmission rate $\beta$ (day$^{-1}$)*

The transmission rate $\beta$ could only be estimated using the data from the transmission experiments. For this analysis detailed information of the infection and transmission process in time is required. This level of detail was not available for the households.

For analysis, each experimental group (mostly pairs, all small scale experiments) of cats was considered as an independent trial and data was collated in the form of the number of Infectious (I), Susceptible (S), and new Cases (C) within a Time interval (d*t*) of one day

(Table S1). These data were analysed using a generalized linear model (GLM) with a binomial error distribution and a cloglog link. Given that $\beta$ is the transmission rate parameter per unit of time $t$, then the probability of new infection (cases) $p$ is

$p = 1 - exp\left(-\beta \frac{I}{N} t\right)$, which upon linearization gives $log(-log(1-p)) = log(\beta) + log(\frac{I}{N}t)$. To fit the GLM $log(\frac{I}{N}t)$ was introduced as an offset variable. The exponent of this model intercept log($\beta$) is the estimated transmission rate parameter $\beta$ *(day*$^{-1}$*).*

***Estimation of the latent and infectious period***

The length of the latent $L$ and infectious period $T$ was quantified by performing a parametric survival analysis where different distributions were assessed. The distributions that best fitted the data (judged by the model with lowest AIC) were an exponential distribution for $L$ and a Weibull distribution for $T$. Because data from the households were left censored, resulting in an uncertain time (mostly > 5 days) of initial exposure of the first infected cat to their infected owners (Table S4), the latent period using household data could not be estimated. The estimation of the infectious period was possible, however, the left censoring for most observations was not considered in the survival model, because the models would not converge. Hence, the estimates are likely an underestimation of the infectious period, when using RT-PCR positive as a correlate of infectiousness.

Tables S3 and S4 show how the data were prepared for this analysis and in Table S5 the estimated parameters of the Weibull distributions of $T$ are provided.

***Estimation of the reproduction number $R_0$***

The reproduction number $R_0$ was estimated as the product of $\beta$ and $T$ (Only for transmission experiments). The *95%* confidence intervals for $R_0$ were derived by Monte Carlo (MC) simulations (1000 replications) assigning to $\beta$ and $T$ lognormal and Weibull distributions respectively. Parameters for these distributions were obtained from the same data set when estimating these parameters as explained above. $R_0$ was also estimated by the final size method (FSM) (6). This method, different from estimating $R_0$ as a results of $\beta*T$ , does not require detailed temporal information of the infection and transmission process. The FSM only uses the information of the number of infected individuals at the end of epidemic, when there is either no more infectious or susceptible individuals in the population. The FSM used for analysis is described in detail elsewhere (7, 8).

Assessment of $R_0 > 1$ was done using the FS method and by MC sampling (when estimating $R_0 = \beta*T$).

*Statistical software*

All analyses were done using the statistical software *R* (9). The library Survival was used for the survival analysis. A detailed description of how to prepare data from a transmission experiment and the R codes for analysis has been reported elsewhere (10).

Table S1. Collated data for the quantification of the transmission rate $\beta$ ($day^{-1}$). Data for each pair of cats (inoculated + contact) was collated daily from day one post inoculation to the day the contact cat was assumed infected (one day before shedding virus).[a]

| Reference | Group | Day | S | I | C | N | dt |
|---|---|---|---|---|---|---|---|
| Halfmann et al.(10) | 1 | 1 | 1 | 1 | 0 | 2 | 1 |
| Halfmann et al.(10) | 1 | 2 | 1 | 1 | 1 | 2 | 1 |
| Halfmann et al.(10) | 2 | 1 | 1 | 1 | 0 | 2 | 1 |
| Halfmann et al.(10) | 2 | 2 | 1 | 1 | 0 | 2 | 1 |
| Halfmann et al.(10) | 2 | 3 | 1 | 1 | 0 | 2 | 1 |
| Halfmann et al.(10) | 2 | 4 | 1 | 1 | 0 | 2 | 1 |
| Halfmann et al.(10) | 2 | 5 | 1 | 1 | 1 | 2 | 1 |
| Halfmann et al.(10) | 3 | 1 | 1 | 1 | 0 | 2 | 1 |
| Halfmann et al.(10) | 3 | 2 | 1 | 1 | 0 | 2 | 1 |
| Halfmann et al.(10) | 3 | 3 | 1 | 1 | 0 | 2 | 1 |
| Halfmann et al.(10) | 3 | 4 | 1 | 1 | 1 | 2 | 1 |
| Gaudreault et al.(12)[b] | 1 | 1 | 1 | 3 | 0 | 4 | 1 |
| Gaudreault et al.(12) | 1 | 2 | 1 | 3 | 1 | 4 | 1 |
| Gaudreault et al.(12) | 2 | 1 | 1 | 3 | 1 | 4 | 1 |
| Bosco-Lauth et al.(11)[c] | 1 | 0 | 2 | 2 | 0 | 4 | 0.5 |
| Bosco-Lauth et al.(11) | 1 | 1 | 2 | 2 | 2 | 4 | 0.5 |
| Bao et al.(14) | P01 | 1 | 1 | 1 | 1 | 2 | 2 |
| Bao et al.(14) | P02 | 1 | 1 | 1 | 1 | 2 | 2 |
| Bao et al.(14) | P03 | 1 | 1 | 1 | 1 | 2 | 2 |
| Bao et al.(14) | P04 | 1 | 1 | 1 | 1 | 2 | 2 |
| Bao et al.(14) | P11 | 1 | 1 | 1 | 0 | 2 | 2 |
| Bao et al.(14) | P12 | 1 | 1 | 1 | 0 | 2 | 2 |
| Bao et al.(14) | P13 | 1 | 1 | 1 | 0 | 2 | 2 |
| Bao et al.(14) | P14 | 1 | 1 | 1 | 0 | 2 | 2 |
| Shi et al. Subadults.(13) | 1 | 1 | 1 | 1 | 1 | 2 | 2 |
| Shi et al. Subadults.(13) | 2 | 1 | 1 | 1 | 0 | 2 | 2 |
| Shi et al. Subadults.(13) | 2 | 2 | 1 | 1 | 0 | 2 | 2 |
| Shi et al. Subadults.(13) | 3 | 1 | 1 | 1 | 0 | 2 | 2 |
| Shi et al. Subadults.(13) | 3 | 2 | 1 | 1 | 0 | 2 | 2 |
| Shi et al. Juveniles.(13) | 1 | 1 | 1 | 1 | 0 | 2 | 1 |
| Shi et al. Juveniles.(13) | 1 | 1 | 1 | 1 | 0 | 2 | 2 |
| Shi et al. Juveniles.(13) | 1 | 3 | 1 | 1 | 1 | 2 | 2 |
| Shi et al. Juveniles.(13) | 2 | 1 | 1 | 1 | 0 | 2 | 1 |
| Shi et al. Juveniles.(13) | 2 | 1 | 1 | 1 | 0 | 2 | 2 |
| Shi et al. Juveniles.(13) | 2 | 3 | 1 | 1 | 0 | 2 | 2 |
| Shi et al. Juveniles.(13) | 2 | 5 | 1 | 1 | 0 | 2 | 2 |
| Shi et al. Juveniles.(13) | 2 | 7 | 1 | 1 | 0 | 2 | 1 |
| Shi et al. Juveniles.(13) | 3 | 1 | 1 | 1 | 0 | 2 | 2 |
| Shi et al. Juveniles.(13) | 3 | 3 | 1 | 1 | 0 | 2 | 2 |
| Shi et al. Juveniles.(13) | 3 | 5 | 1 | 1 | 0 | 2 | 2 |
| Shi et al. Juveniles.(13) | 3 | 7 | 1 | 1 | 0 | 2 | 1 |

[a] Group = identifies the experiment group, Day = day post inoculation, S = number of Susceptible at the start of the day, I = number of infectious, C = number of new infections “cases” confirmed the next day, N = total number of animals in the experiment, dt = time interval in days.

[b] Samples were taken every two days. And all contacts were already positive two days post challenge. When using a dt = 2 (only one row per group), it was not possible to estimate $\beta$. Hence to be able to analyse this data, we had to assume that one contact in for instance group 1 became infected at day 2 and the other contact in group 2 at day 1. In this way we used a  dt = 1.

[c] Both contacts were infected the next day post challenge. By using a dt = 1 (only one row), it was not possible to estimate $\beta$. Hence, to be able to make estimations, we divided the day in two periods (dt = 0.5) and assumed transmission took place during the second period.

Table S2. Collated data for the estimation of the infectious and latent periods.[a]

| Reference | Treatment | Shedding | Time | Time2 | Event | LPtime | LPtime2 | Event_Lp | Peak shedding | Units |
|---|---|---|---|---|---|---|---|---|---|---|
| Halfmann et al.(10) | Inoculated | nasal | 6 | 6 | 1 | 1 | 1 | 1 | 3.6 | PFU |
| Halfmann et al.(10) | Inoculated | nasal | 4 | 4 | 1 | 2 | 2 | 2 | 2.8 | PFU |
| Halfmann et al.(10) | Inoculated | nasal | 6 | 6 | 1 | 1 | 1 | 1 | 4 | PFU |
| Halfmann et al.(10) | Contact | nasal | 5 | 5 | 1 | 1 | 3 | 3 | 4.5 | PFU |
| Halfmann et al.(10) | Contact | nasal | 4 | 4 | 1 | 5 | 5 | 2 | 4.1 | PFU |
| Halfmann et al.(10) | Contact | nasal | 5 | 5 | 1 | 4 | 4 | 2 | 3.5 | PFU |
| Bosco-Lauth et al.(11) | Inoculated | nasal | 4 | 4 | 1 | 1 | 1 | 1 | 3.9 | PFU |
| Bosco-Lauth et al.(11) | Inoculated | nasal | 4 | 4 | 1 | 1 | 1 | 1 | 4.3 | PFU |
| Bosco-Lauth et al.(11) | Inoculated | nasal | 4 | 4 | 1 | 1 | 1 | 1 | 3.7 | PFU |
| Bosco-Lauth et al.(11) | Inoculated | nasal | 5 | 5 | 0 | 1 | 1 | 1 | 6.3 | PFU |
| Bosco-Lauth et al.(11) | Inoculated | nasal | 5 | 5 | 0 | 1 | 1 | 1 | 2.3 | PFU |
| Bosco-Lauth et al.(11) | Contact | nasal | 6 | 8 | 3 | 1 | 1 | 1 | 3.6 | PFU |
| Bosco-Lauth et al.(11) | Contact | nasal | 6 | 8 | 3 | 1 | 1 | 1 | 4.4 | PFU |
| Gaudreault et al.(12) | Contact | nasal | 7 | 11 | 3 | 1 | 3 | 3 | 9 | RNA |
| Gaudreault et al.(12) | Contact | nasal | 4 | 4 | 1 | 1 | 3 | 3 | 9 | RNA |
| Shi et al. Subadults.(13) | Inoculated | faeces | 5 | 7 | 3 | 3 | 3 | 2 | 5.3 | RNA |
| Shi et al. Subadults.(13) | Inoculated | faeces | 5 | 7 | 3 | 3 | 3 | 2 | 4.9 | RNA |
| Shi et al. Subadults.(13) | Inoculated | faeces | 5 | 7 | 3 | 3 | 3 | 2 | 4.5 | RNA |
| Shi et al. Juveniles.(13) | Inoculated | nasal | 6 | 8 | 3 | 2 | 2 | 2 | 7.5 | RNA |
| Shi et al. Juveniles.(13) | Inoculated | nasal | 8 | 10 | 3 | 2 | 2 | 2 | 7.5 | RNA |
| Shi et al. Juveniles.(13) | Inoculated | nasal | 8 | 10 | 3 | 2 | 2 | 2 | 7.9 | RNA |
| Bao et al. (14) | Inoculated | nasal | 11 | 11 | 1 | 1 | 1 | 1 | 4.5 | RNA |
| Bao et al. (14) | Inoculated | nasal | 11 | 13 | 3 | 1 | 1 | 1 | 5.6 | RNA |
| Bao et al. (14) | Inoculated | nasal | 11 | 13 | 3 | 1 | 1 | 1 | 4.3 | RNA |
| Bao et al. (14) | Inoculated | nasal | 13 | 13 | 1 | 1 | 1 | 1 | 5.2 | RNA |
| Bao et al. (14) | Contact | nasal | 9 | 11 | 3 | 2 | 2 | 2 | 4 | RNA |
| Bao et al. (14) | Contact | nasal | 11 | 13 | 3 | 2 | 2 | 2 | 3.7 | RNA |
| Bao et al. (14) | Contact | nasal | 5 | 7 | 3 | 2 | 2 | 2 | 2.8 | RNA |

| | | | | | | | | | | |
|---|---|---|---|---|---|---|---|---|---|---|
| Bao et al. (14) | Contact | nasal | 9 | 11 | 3 | 2 | 2 | 2 | 3.1 | RNA |

[a] Each row represents observations for an individual cat. Time = number of days the cat was assumed infectious (shedding virus) and detected positive for the last time; Time2 = time in days when cat was first negative; event = categorical variable where 1 = cat recovered (shedding stopped), 0 = cat was still shedding at the end of the experiment (right censoring) or 3 = shedding stopped between time and time2. LPtime, LPtime2 and LPevent are the data used for the estimation of the latent period. LPtime = day animal last detected negative (starts shedding), LPtime2 = animal start shedding, LPevent = categorical variable where 1 = event observed at time 1, 2 = left censoring, 3 = shedding started between LPtime and LPtime2.

Table S3. Collated data from infected households with more than one cat. These data were used for the estimation of the reproductive number $R_0$ using the final size method.

| Study | Household id | No. of humans | No. of cats | PCR+ cats | Serology + cats[a] | Last day serum sample[b] |
|---|---|---|---|---|---|---|
| Chaintoutis et al.(4) | 1 | 1 | 3 | 2 | 1 | 63 |
| Hamer et al.(7) | D | NP[c] | 2 | 2 | 2 | 93 |
| Hamer et al.(7) | OO | NP | 3 | 3 | 3 | 38 |
| Klaus et al.(5) | 2 | 2 | 2 | 1 | 1 | 36 |
| Segales et al.(9) | 3 | 1 | 2 | 2 | 2 | 10 |
| Neira et al.(6) | 1 | NP | 2 | 1 | 1 | 39 |
| Neira et al.(6) | 2 | 2 | 3 | 3 | 1 | 40 |
| Goryoka et al.(8) | 2 | 3 | 2 | 2 | 2 | 20 |

[a] Only cats serology positive (+) were considered infected for estimation of $R_0$.

[b] This is the time from the day of the first cat or human confirmation of infection in the household and the time the last serum sample for serology was taken from all cats in the household. These data help to confirm the assumption that the transmission process within each household reached its end (no more infectious or susceptible cats present in the household).

[c] NP = not provided

Table S4. Collated data from observational studies describing the longitudinal follow up of infection in infected cats from infected households. These data were used to estimate the duration of observed shedding in naturally infected cats.[a]

| Study | Houshold ID | Cat ID | Sample | Cat's Age (Years) | Time | Time2 | Event | Owner symptoms (Days)[a] | Maximum observed shedding | Units |
|---|---|---|---|---|---|---|---|---|---|---|
| Chaintoutis et al.(4) | 1 | C2 | Oropharyngeal | 10 | 7 | 7 | 1 | -7 | 7 | $log_{10}$RNAcopies/swab |
| Chaintoutis et al.(4) | 1 | C3 | Oropharyngeal | 10 | 7 | 7 | 1 | -7 | 8.5 | $log_{10}$RNAcopies/swab |
| Chaintoutis et al.(4) | 1 | C2 | Fecal | 10 | 5 | 5 | 1 | -7 | 5.6 | $log_{10}$RNAcopies/swab |
| Chaintoutis et al.(4) | 1 | C3 | Fecal | 10 | 7 | 7 | 1 | -7 | 6.6 | $log_{10}$RNAcopies/swab |
| Barrs et al.(15) | 1 | DHS/7/F | Nasal | 7 | 11 | 11 | 1 | -10, -1 | 6.3 (21.3)[d] | $log_{10}$RNAcopies/swab |
| Barrs et al.(15) | 1 | DHS/7/F | Oral | 7 | 7 | 9 | 3 | -10, -1 | 5.6 (22.9) | $log_{10}$RNAcopies/swab |
| Barrs et al.(15) | 1 | DHS/7/F | Rectal | 7 | 1 | 1 | 1 | -10, -1 | 3.2 (33) | $log_{10}$RNAcopies/swab |
| Barrs et al.(15) | 2 | AHS/13/M | Oral | 13 | 4 | 9 | 3 | NP[c] | 22 | Ct[c] |
| Barrs et al.(15) | 3 | SSH/5/M | Oral | 5 | 5 | 10 | 3 | NP | 26.8 | Ct |
| Neira et al.(15) | 1 | Cat 1 | Nasal | 10 | 5 | 5 | 1 | -7, -6 | 31 | Ct |
| Neira et al.(15) | 1 | Cat 1 | Fecal | 10 | 4 | 4 | 1 | -7, -6 | 30.8 | Ct |
| Neira et al.(15) | 1 | Cat 2 | Nasal | 10 | 17 | 17 | 1 | -7, -6 | 21.9 | Ct |
| Neira et al.(15) | 1 | Cat 2 | Fecal | 10 | 9 | 9 | 1 | -7, -6 | 31.2 | Ct |
| Neira et al.(15) | 1 | Cat 3 | Nasal | 10 | 7 | 10 | 3 | -7, -6 | 29.9 | Ct |
| Neira et al.(15) | 1 | Cat 3 | Nasal | 10 | 7 | 7 | 1 | -7, -6 | 29.9 | Ct |
| Bessiere et al.(16) | 3 | 1 | Oropharyngeal | 3 | 2 | 2 | 1 | -5 | 35.6 | Ct |

[a] For the estimation of the duration of shedding, each row represents observations for an individual cat within a household. The following columns were used: Time = number of days the cat detected RT-PCR positive and detected positive for the last time; Time2 = time in days when cat was first negative; Event = categorical variable where 1 = cat recovered (shedding stopped, no censored data), 0 = cat was still shedding at the end of the experiment (right censoring) or 3 = shedding stopped between Time and Time2.

[b] This is number of days between the onset of clinical signs in the cat's owner(s) (multiple values, represent multiple people in the household) and the first detection of infection in the cat. A value of -7 means the owner first had clinical signs 7 days before positive confirmation of infection in the cat. For

estimation of the duration of shedding this left censoring was ignored and duration of shedding was considered from the first day of detection of infection until the last day that the cat was detected positive by RT-PCR. Hence Estimates of duration of shedding (as measured by PCR) are likely to be underestimated.

[c] Ct = RT-PCR cycle threshold. NP = data not provided.

[d] Data in brackets are the RT-PCR Ct values.

Table S5. Estimated Weibull parameters (Shape and Scale) describing the length of the infectious period *T*.

| Study | Infection route | Shape | Scale |
|---|---|---|---|
| *Direct transmission* | | | |
| Halfmann et al.(10) | Contact | 7.8 | 4.8 |
| | Inoculation | 7.8 | 5.7 |
| Bosco-Lauth et al.(11) | Contact | 7.8 | 7.1 |
| | Inoculated | 7.8 | 4.9 |
| Gaudreault et al.(12) | Contact | 7.8 | 7.0 |
| Bao et al.(14) | Contact | 7.8 | 10.5 |
| | Inoculated | 7.8 | 12.2 |
| *Droplet transmission* | | | |
| Shi et al. juveniles.(13) | Inoculated | 6.6 | 8.2 |
| Shi et al. subadults.(13) | Inoculated | 6.6 | 5.8 |
| *Observational studies* | | | |
| Household cats (4,6,15,16) | Contact | 2.0 | 7.9 |
